# IPC - Isoelectric Point Calculator

**DOI:** 10.1101/049841

**Authors:** Lukasz P. Kozlowski

## Abstract

Accurate estimation of the isoelectric point (*pI*) based on the amino acid sequence can be useful for many biochemistry and proteomics techniques such as 2-D polyacrylamide gel electrophoresis, or capillary isoelectric focusing used in combination with high-throughput mass spectrometry. Here, I present the Isoelectric Point Calculator, a web service for the estimation of *pI* using different sets of dissociation constant (*pKa*) values, including two new, computationally optimized *pKa* sets. According to the presented benchmarks, IPC outperform previous algorithms by at least 14.9% for proteins and 0.9% for peptides (on average, 22.1% and 59.6%, respectively), which corresponds to an average error of the *pI* estimation equal to 0.87 and 0.25 pH units for proteins and peptides, respectively. Peptide and protein datasets used in the study and the precalculated *pI* for PDB, SwissProt databases are available for large-scale analysis and future development. The IPC can be accessed at http://isoelectric.ovh.org.

## Introduction

Analysis of proteins starts from the heterogeneous mixture (lysate) from which protein fraction needs to be isolated. Next, individual proteins are separated and finally identified. The procedure relies on physicochemical properties of amino acids such as a molecular mass or a charge. Over the years, many techniques were introduced to allow to accomplish the task. One of the oldest, but still widely used technique is 2-D polyacrylamide gel electrophoresis (2D-PAGE)^1,2^, where proteins are separated in two dimensions on a gel and identified using estimated molecular weight and isoelectric point (*pI* is the pH value at which the net charge of a macromolecule is zero, and therefore its electrophoretic mobility is stopped). Unfortunately, 2D-PAGE suffers from several intrinsic technical problems (e.g., performs poorly for very large, very small, extremely acidic or basic proteins). Therefore, 2D-PAGE has been today replaced in many cases by gel-free techniques such as high-throughput mass spectrometry (MS)^3,4^. Before the mass spectrometry is applied, the sample is digested by trypsin into short peptides and then fractionated by isoelectric focusing into so called fractions which allows to reduce MS analysis complexity. Although molecular techniques for protein analysis have changed, the interpretation of the results from those techniques in many cases rely on accurate estimations of *pI* for reference polypeptides.

For polypeptides, *pI* depends mostly on the acid dissociation constants (*pKa*) of the ionizable groups of seven charged amino acids: glutamate (δ-carboxyl group), aspartate (ß-carboxyl group), cysteine (thiol group), tyrosine (phenol group), histidine (imidazole side chains), lysine (ε-ammonium group) and arginine (guanidinium group). Additionally, the charge of the amine and carboxyl terminal groups contribute to *pI* and can greatly affect *pI* of short peptides^5^. Overall, the net charge of the protein or peptide is strongly related to the solution (buffer) pH and can be approximated using the Henderson-Hasselbalch equation^6^. It should be kept in mind that the values of dissociation constants used in the calculations are usually derived empirically and can vary substantially depending on the experimental setup such as temperature or buffer ionic strength (herein presented method, Isoelectric Point Calculator, is compared to 15 such *pKa* sets). On the other hand, *pKa* values or *pI* can be derived computationally giving the large sets of proteins or peptides for which *pI* information is known. This is the approach, presented in this study. The problem of computational prediction of *pI* was already addressed by two other research groups using artificial neural networks (ANN)^7^ and support vector machines (SVM)^8,9^. Here, I present IPC program which is based on the optimization using a Basin-Hopping procedure^10^. Presented results shows that IPC overperform all currently, available algorithms.

## Results

### Comparison to other algorithms

To compare the performance of Isoelectric Point Calculator fifteen, other *pKa* sets and two programs based on SVM (pIR) and ANN (pIPredict) were selected. Isoelectric point predictions were validated separately for peptides and proteins as they differ substantially. Proteins are relatively big molecules with a plethora of charged residues. Moreover, in the proteins *pI* is affected by many, additional factors such as post translational modifications, solvent accessibility, etc. On the other hand, peptides are short, possessing usually only a handful of charged residues and therefore their *pI* is easier to predict. In the presented study two protein databases, SWISS-2DPAGE and PIP-DB, were used. For peptides, three datasets from separate high-throughput experiments were used. At the beginning, two databases for proteins were merged. As the content of the databases overlapped and was redundant, additional post processing and cleaning of the data was necessary. First of all, not all records contained useful information, namely isoelectric point and sequence or Uniprot ID. Moreover, even separate databases were redundant (contained multiple records with the same sequence or Uniprot ID). Therefore, the duplicates were merged into unique records and *pI* information was averaged if needed (multiple *pI* values coming from separate experiments). Next, the worst outliers defined here as those proteins for which the difference between the experimental *pI* and the average predicted *pI* was greater than the threshold of the mean standard error (MSE) of 3 were excluded as they represented possible annotation errors. Finally, the resulting dataset consisting of over 2,000 proteins was divided into a training set (75% randomly chosen proteins) and a testing set. The training set was used to obtain optimized *pKa* values and the test set was used to evaluate IPC on proteins not used during training. A similar procedure was employed to peptide datasets with the exception that then the threshold of MSE of 0.25 was used (for more details see Methods). The results of the benchmarks for *pI* prediction are presented in Table 1–3. Table 1 shows the results on testing sets both for proteins and peptides. IPC produced best results (the lowest RMSD and the smallest number of outliers). For comparison the results on the training set are presented in Table 2. The performance of the IPC_protein set is slightly better for the 75% training dataset (RMSD of 0.8376 for the 75% training set versus 0.8731 for the 25% test set), but this is expected (even though optimization procedure was cross validated the overfitting cannot be avoided fully, but results in Table 1 and Table 2 show that this is not critical in this case). Moreover, the general performance of IPC do not depend on the datasets used for training (Table 3). Furthermore, the results for the training sets and the results for the test sets are consistent (Table 1 and Table 2, respectively).In most cases the order of the method’s performance on both training and testing datasets is similar; for instance the change in the order on the protein dataset can be seen for the Dawson and Bjellqvist *pKa* sets, which is within the error margin. Similarly, there are some changes in the method order depending on the peptide dataset, but only for methods with a very similar performance, e.g., Lehninger and Solomon on PIP-DB. In most cases, the change is within the margin of error. The IPC sets, regardless of the dataset and the validation procedures, performed the best. Similar results are obtained when comparing the number of outliers produced by the individual *pKa* sets. Outliers correspond to cases of extremely poor prediction (the difference between the predicted and experimental pI is greater than an arbitrarily chosen threshold; e.g., for proteins, an MSE of 3 was used as the threshold). In all cases, IPC produced the smallest number of outliers. It should be stressed, that all algorithms, except IPC, pIR and pIPredict, rely on experimentally derived *pKa* values and therefore they were not optimized for particular data sets. As IPC results were validated on test set not used in training, the only remaining algorithms which may be optimized towards a particular dataset are pIR and pIPredict. pIR is a support vector machine method which used PIP-DB proteins for training, thus it is interesting to investigate how it performs on the other protein set. As one can see in Table 3, while pIR produce reasonable results for the PIP-DB dataset, its predictive performance decreases significantly on the SWISS-2DPAGE dataset. This means that pIR method was most likely overfitted towards PIP-DB proteins (move from the middle of the table - PIP-DB dataset, to the bottom - SWISS-2DPAGE dataset). Moreover, it should be stressed that all benchmarks from Audain et al. and presented here describing PIP-DB cannot be compared directly as they were done on different subsets of PIP-DB (Audain et al. removed all records which have more than one *pI* measurement for given protein, while here average was used instead). Also, pIPredict performs worse than most of the methods. Most likely it is due the fact that pIPredict was trained only on peptide dataset from Gauci et al., which is smaller than used in the presented study. Moreover, it was not trained on any protein dataset, thus pIPredict should rather be used only for peptides.

**Table 1.** Prediction of isoelectric point on the 25% testing datasets

| Protein dataset |  |  |  | Peptide dataset |  |  |  |
| --- | --- | --- | --- | --- | --- | --- | --- |
| Method | RMSD | % | Outliers | Method | RMSD | % | Outliers |
| IPC_protein | 0.874 | 0 | 46 | IPC_peptide | 0.251 | 0 | 232 |
| Toseland | 0.934 | 14.9 | 52 | Solomon | 0.255 | 0.9 | 235 |
| Bjellqvist | 0.944 | 17.7 | 47 | Lehninger | 0.262 | 2.5 | 236 |
| Dawson | 0.945 | 17.8 | 56 | EMBOSS | 0.325 | 18.5 | 372 |
| Wikipedia | 0.955 | 20.5 | 55 | Wikipedia | 0.421 | 47.9 | 1467 |
| Rodwell | 0.963 | 22.8 | 58 | Toseland | 0.425 | 49.1 | 990 |
| ProMoST | 0.966 | 23.6 | 52 | Sillero | 0.428 | 50.3 | 1223 |
| Grimsley | 0.968 | 24.2 | 60 | Dawson | 0.435 | 52.9 | 1432 |
| Solomon | 0.970 | 24.8 | 58 | Thurkill | 0.481 | 69.7 | 1361 |
| Lehninger | 0.970 | 25.0 | 59 | Rodwell | 0.502 | 78.4 | 1359 |
| pIR | 1.013 | 38.0 | 58 | DTASelect | 0.550 | 99.1 | 1714 |
| Nozaki | 1.024 | 41.3 | 56 | Nozaki | 0.602 | 124.3 | 1368 |
| Thurkill | 1.030 | 43.4 | 61 | Grimsley | 0.616 | 131.4 | 1550 |
| DTASelect | 1.032 | 44.1 | 58 | Bjellqvist | 0.669 | 161.5 | 1583 |
| pIPredict | 1.048 | 49.4 | 56 | pIPredict | 1.024 | 493.6 | 2720 |
| EMBOSS | 1.056 | 52.3 | 69 | ProMoST | 1.239 | 873.4 | 2649 |
| Sillero | 1.059 | 53.2 | 63 | pIR | 1.881 | 4159.7 | 3358 |
| Patrickios | 2.392 | 3201.8 | 227 | Patrickios | 1.998 | 5479.1 | 2739 |
| Avg_pl* | 0.960 | 22.1 | 53 | Avg_pl | 0.454 | 59.6 | 1571 |

\* Average from all *pKa* sets without Patrickios (highly simplified *pKa* set) and IPC sets. Note, that the average *pI* is calculated on the level of individual protein or peptide, thus it does not represent the average from values presented in the table for individual methods
% - Note that the pH scale is logarithmic with base 10; thus, the percent difference corresponds to $\text{pow}(10, x)$ , where $x$ is equal to the delta of the RMSD of two error estimates represented in pH units; for example, the % difference between Toseland and IPC\_protein is $\text{pow}(10, (0.934-0.874))$

**Table 2.** Prediction of isoelectric point on the 75% training datasets

| Protein dataset |  |  |  | Peptide dataset |  |  |  |
| --- | --- | --- | --- | --- | --- | --- | --- |
| Method | RMSD | % | Outliers | Method | RMSD | % | Outliers |
| IPC_protein | 0.838 | 0 | 114 | IPC_peptide | 0.247 | 0 | 635 |
| Toseland | 0.898 | 15.0 | 131 | Solomon | 0.251 | 0.8 | 638 |
| **Bjellqvist** | **0.922** | **21.5** | **149** | Lehninger | 0.256 | 2.4 | 643 |
| **Dawson** | **0.920** | **20.9** | **156** | EMBOSS | 0.322 | 18.8 | 1088 |
| Wikipedia | 0.930 | 23.8 | 157 | Wikipedia | 0.413 | 46.3 | 4280 |
| Rodwell | 0.938 | 26.1 | 159 | Sillero | 0.426 | 50.9 | 3025 |
| ProMoST | 0.938 | 26.1 | 140 | Toseland | 0.427 | 51.2 | 3618 |
| Grimsley | 0.939 | 26.2 | 147 | Dawson | 0.432 | 52.9 | 4192 |
| **Solomon** | **0.947** | **28.5** | **159** | Thurkill | 0.480 | 70.8 | 4017 |
| **Lehninger** | **0.947** | **28.7** | **160** | Rodwell | 0.506 | 81.2 | 4061 |
| **pIR** | **1.026** | **54.2** | **180** | DTASelect | 0.541 | 96.8 | 4902 |
| **Nozaki** | **1.005** | **47.1** | **169** | Nozaki | 0.599 | 124.8 | 4013 |
| **Thurkill** | **1.018** | **51.5** | **173** | Grimsley | 0.611 | 130.9 | 4609 |
| **DTASelect** | **1.017** | **51.1** | **167** | Bjellqvist | 0.661 | 159.2 | 4672 |
| **pIPredict** | **1.057** | **65.9** | **173** | pIPredict | 1.024 | 497.8 | 8051 |
| EMBOSS | 1.040 | 59.4 | 189 | ProMOST | 1.233 | 867.5 | 7999 |
| Sillero | 1.042 | 60.1 | 188 | pIR | 1.862 | 4020.9 | 9921 |
| Patrickios | 2.237 | 2405.1 | 645 | Patrickios | 1.977 | 5266.8 | 8131 |
| Avg_pl* | 0.940 | 26.6 | 151 | Avg_pl | 0.451 | 59.7 | 4600 |

\* Average from all *pKa* sets without the Patrickios (highly simplified *pKa* set) and IPC sets. Note, that the average *pI* is calculated on the level of individual protein or peptide
Outliers correspond to the number of predictions for which the difference between the experimental *pI* and the predicted *pI* exceeded the threshold of an MSE of 3 for the protein dataset and an MSE of 0.25 for the peptide dataset.

**Table 3.**
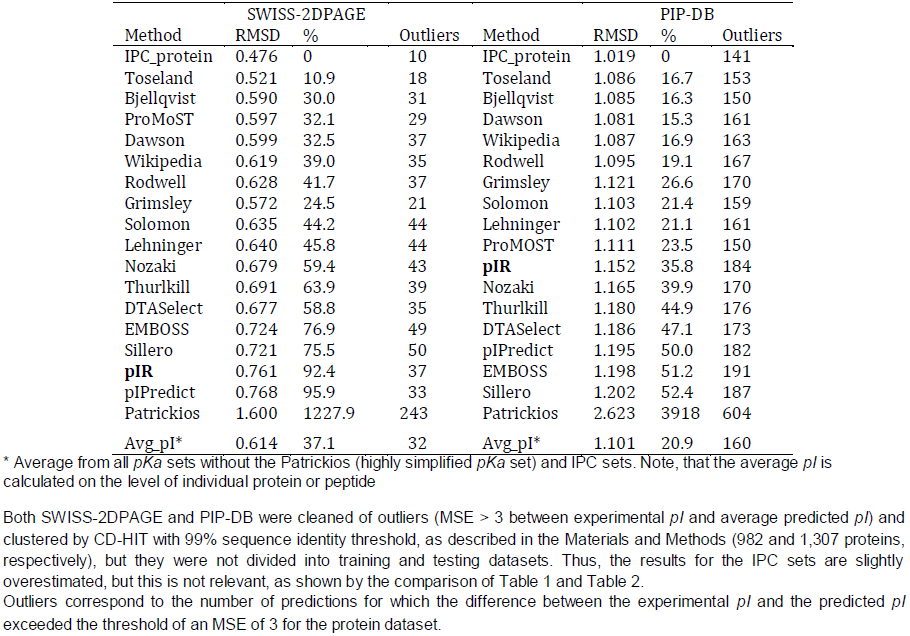
Detailed statistics for the different *pKa* sets for SWISS-2DPAGE and PIP-DB

| SWISS-2DPAGE |  |  |  | PIP-DB |  |  |  |
| --- | --- | --- | --- | --- | --- | --- | --- |
| Method | RMSD | % | Outliers | Method | RMSD | % | Outliers |
| IPC_protein | 0.476 | 0 | 10 | IPC_protein | 1.019 | 0 | 141 |
| Toseland | 0.521 | 10.9 | 18 | Toseland | 1.086 | 16.7 | 153 |
| Bjellqvist | 0.590 | 30.0 | 31 | Bjellqvist | 1.085 | 16.3 | 150 |
| ProMoST | 0.597 | 32.1 | 29 | Dawson | 1.081 | 15.3 | 161 |
| Dawson | 0.599 | 32.5 | 37 | Wikipedia | 1.087 | 16.9 | 163 |
| Wikipedia | 0.619 | 39.0 | 35 | Rodwell | 1.095 | 19.1 | 167 |
| Rodwell | 0.628 | 41.7 | 37 | Grimsley | 1.121 | 26.6 | 170 |
| Grimsley | 0.572 | 24.5 | 21 | Solomon | 1.103 | 21.4 | 159 |
| Solomon | 0.635 | 44.2 | 44 | Lehninger | 1.102 | 21.1 | 161 |
| Lehninger | 0.640 | 45.8 | 44 | ProMOST | 1.111 | 23.5 | 150 |
| Nozaki | 0.679 | 59.4 | 43 | **pIR** | 1.152 | 35.8 | 184 |
| Thurlkill | 0.691 | 63.9 | 39 | Nozaki | 1.165 | 39.9 | 170 |
| DTASelect | 0.677 | 58.8 | 35 | Thurlkill | 1.180 | 44.9 | 176 |
| EMBOSS | 0.724 | 76.9 | 49 | DTASelect | 1.186 | 47.1 | 173 |
| Sillero | 0.721 | 75.5 | 50 | pIPredict | 1.195 | 50.0 | 182 |
| **pIR** | 0.761 | 92.4 | 37 | EMBOSS | 1.198 | 51.2 | 191 |
| pIPredict | 0.768 | 95.9 | 33 | Sillero | 1.202 | 52.4 | 187 |
| Patrickios | 1.600 | 1227.9 | 243 | Patrickios | 2.623 | 3918 | 604 |
| Avg_pI* | 0.614 | 37.1 | 32 | Avg_pI* | 1.101 | 20.9 | 160 |

\* Average from all *pKa* sets without the Patrickios (highly simplified *pKa* set) and IPC sets. Note, that the average *pI* is calculated on the level of individual protein or peptide
Outliers correspond to the number of predictions for which the difference between the experimental *pI* and the predicted *pI* exceeded the threshold of an MSE of 3 for the protein dataset.

## Auxiliary statistics

Figures 1 and Figures 2 show the correlation plots between the experimental and theoretical isoelectric points for proteins and peptides on different datasets calculated using different *pKa* sets. These plots are useful to assess the quality of the datasets used. The Pearson correlations (R^2^) between a *pKa* set, e.g., EMBOSS, give a good impression of the quality of the dataset and the number of outliers, which were defined here as those where the MSE exceeded 3 for the average pI prediction (this corresponds to ~1.73 pH unit difference). Even if we assume that the presented, nine parametric model is highly simplified e.g., it does not take post-translational modifications into account, we can suspect that such a large difference is more likely an annotation error in the database than a true difference (this assumption was confirmed by randomly checking some outliers; data not shown, available on request). Moreover, contrary to previous works, R^2^ was not used as a performance measure because it should not be considered in this way. R^2^ measures how well the current model fits a linear model. It is unlikely that the experimental isoelectric point can be explained using a highly simplified nine parametric model that does not take into account multiple factors (see Methods for more details). The R^2^ value is a useful statistic for preliminary analysis but should not be used for evaluating the performance. Similarly, scatter plots between the experimental *pI* and those produced by different *pKa* sets (Fig. 2) can give a good impression of the correctness of the model, but quantitative measurement of the performance requires better measures, e.g., the root-mean-square deviation (RMSD), which presents the sample standard deviation of the differences between the predicted values and the observed values. An additional advantage of the RMSD is that it is simple to explain and reflects the error of the prediction in pH units. Another performance metric used here is the number of outliers at a given threshold (for the protein dataset the threshold was set to MSE > 3 between the experimental *pI* and average prediction *pI* for removing outliers from the datasets; in this way, none of the *pKa* sets was favored). For instance, the Patrickios *pKa* set is highly simplified and generally should not be used. Thus, this set was not included in the average calculation. In all benchmarks, the Patrickios*pKa* set performed the worst. As illustrated in Fig. 2 (top, right panel), this set cannot correctly predict the *pI* for proteins with *pI* > 6, but it performs relatively well in the 4-6 *pI* range.

**Figure 1.**
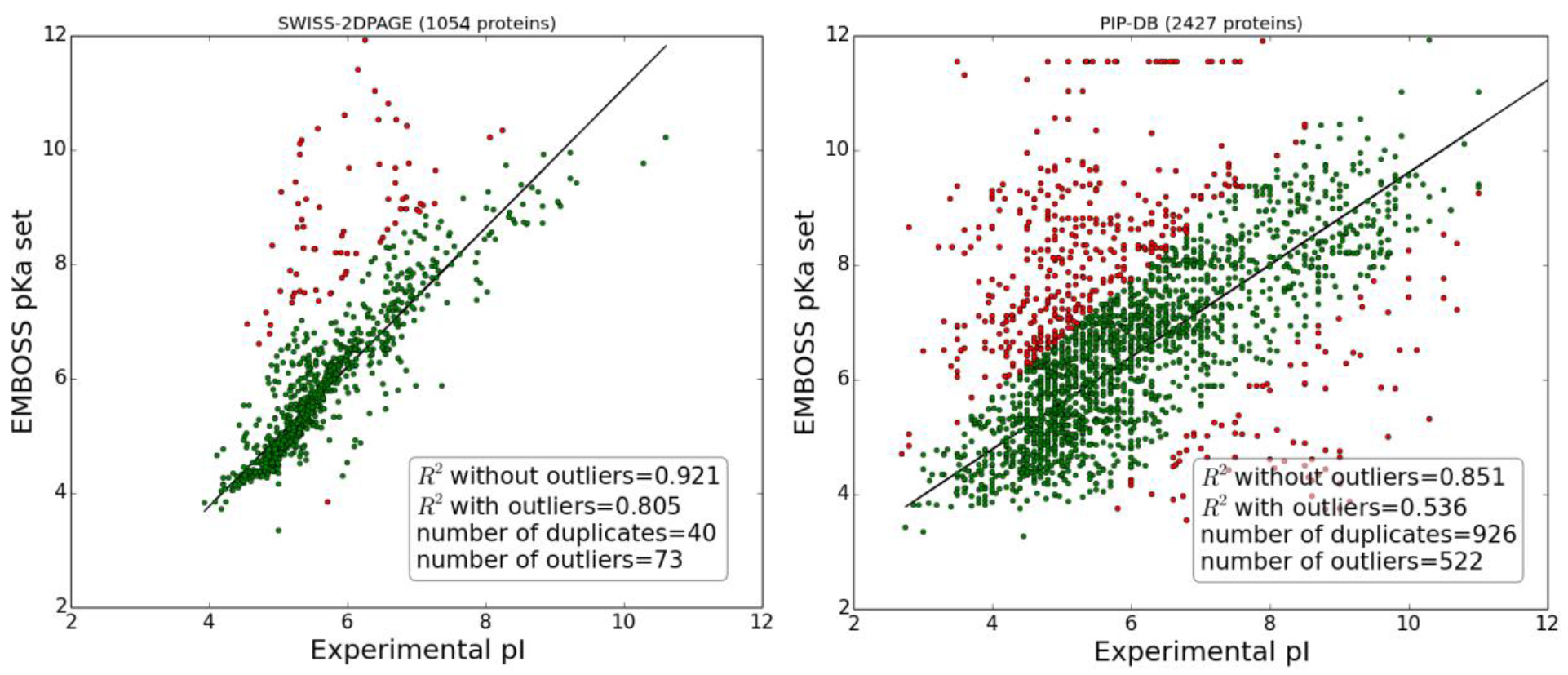
Correlation plots of the experimental versus the theoretical isoelectric points for protein datasets (SWISS-2DPAGE and PIP-DB) calculated using the EMBOSS *pKa* set. Outliers are defined as MSE > 3 and are marked in red. Plots correspond to datasets as presented by the authors before cleaning and the removal of duplicates (duplicates are defined as records that have the same sequence but are referred to as separate records in the database). In both databases, the authors report multiple *pI* values from different experiments for the same sequences in separate records. For the current analysis, the average *pI* was used. The solid line represents the linear regression after removal of the outliers.

**Figure 2.**
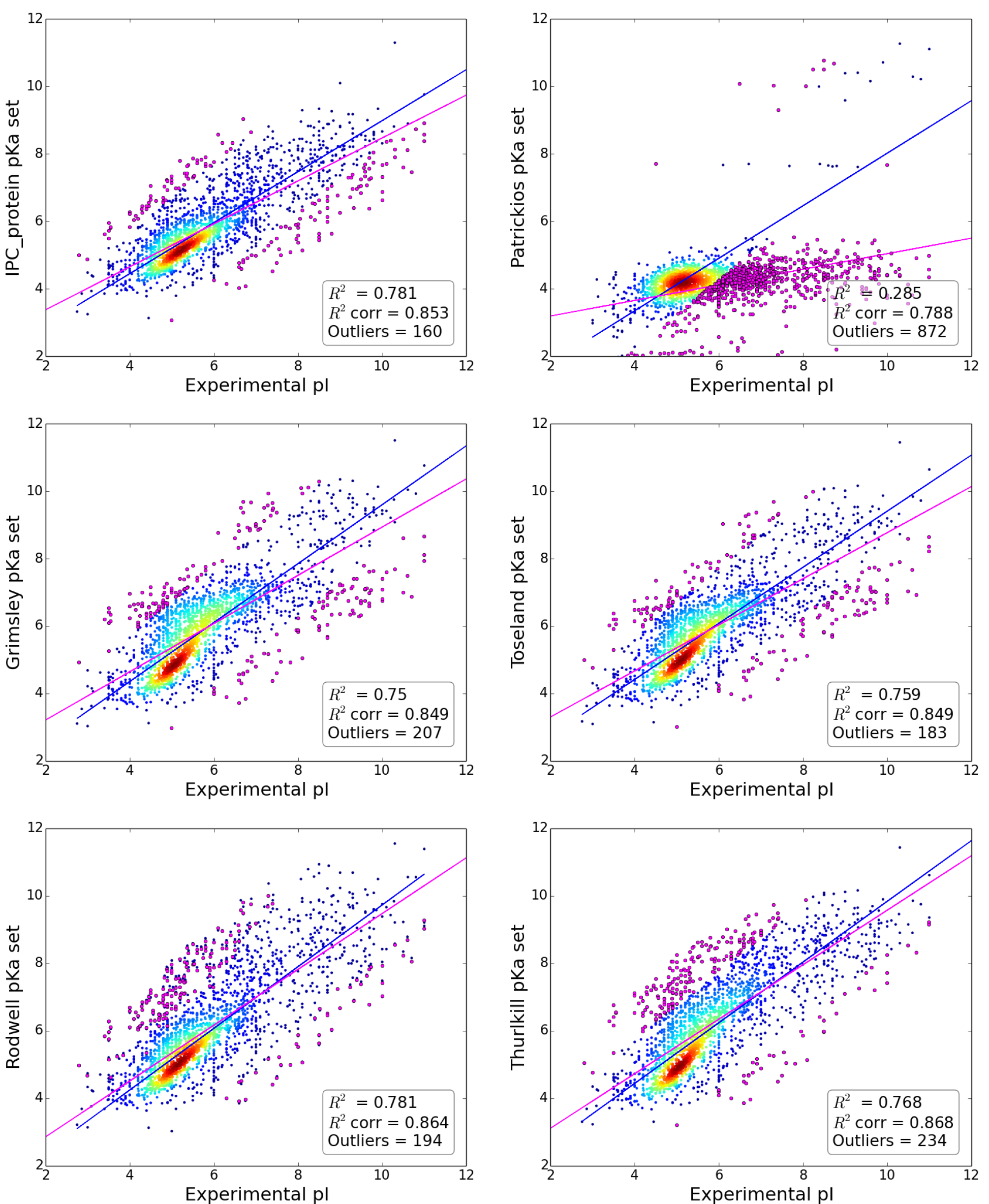
Correlation plots of the experimental versus theoretical isoelectric points for the main protein dataset (merge of SWISS-2DPAGE and PIP-DB, including the training and test sets) calculated using different *pKa* sets.R^2^corr - Pearson correlation after the removal of outliers.R^2^corr-Pearson correlation after the removal of outliers. Additionally, the linear regression models fitted to predictions with outliers (magenta line) and without outliers (blue line) are shown. Outliers (marked in magenta) are defined as *pI* predictions with MSE > 3 in comparison to the experimental *pI*. Other predictions are represented as heat maps according to the density of points. The numbers of outliers for both the training and testing set are shown together. For brevity, only six *pKa* sets are shown.

## Discussion

The distribution of the isoelectric points of proteins in proteomes is universal for almost all organisms^11^, which can be demonstrated by plotting isoelectric points of the proteins stored in the *SwissProt* database. The distribution is bimodal with a low fraction of proteins with a *pI* close to 7.4. This is because the proteins are mostly insoluble, less reactive and unstable at pH close to their *pI*. The pH inside of most cells is close to 7.4, therefore this property of proteomes can be a result of evolutionary selection or simply a result of the chemical properties of amino acids^12^. Naturally, there are some exceptions. Some halophilic Archaea organisms do not try to fight the high concentration of salt in their environment; instead, they change the physiological pH inside their cells to be more similar to the environment (in this way, they use less energy to maintain homeostasis)^13^. This response has dramatic consequences for the amino acid compositions and isoelectric points of their proteins (Fig. 3).

It should be stressed that the relative difference between the different *pKa* sets is often small and statistically insignificant (e.g., *pI* calculated by Bjellqvist vs. Dawson *pKa* sets on protein datasets), but even general knowledge of which *pKa* sets are better and which should be used for a particular type of data (e.g., protein versus peptides) is not commonly used (Fig. 3, bottom two panels). Furthermore, presented results demonstrate that prediction of *pI* is easier for short peptides than for proteins as the former contain less charged and modified amino acids (e.g. compare RMSD values between peptide and protein datasets). Similarly, the dataset on which methods are trained and/or evaluated can result in different estimations of RMSD error. For example, Fig. 1 shows that PIP-DB contains multiple outliers and duplicates in comparison to SWISS-2DPAGE. This noise in the data leads to almost a doubling of the RMSD (Table 3).Nevertheless, the method order is usually preserved.

As mentioned earlier, one of the main limitations of IPC is that it uses a nine parametric model which is a highly simplistic approximation, and do not take into account many aspects of proteins such as post translational modification. It should be noted that posttranslational modifications occur much more frequently in Eukaryotic proteins than in Prokaryotic, thus it is interesting to investigate how accurately *pI* can be predicted in these two kingdoms separately. As illustrated in Supplementary Table 1 all *pI* prediction methods perform better on prokaryotic proteins. This proves that when working with Eukaryotic proteins one should keep in mind that *pI* prediction accuracy can be decreased due possible posttranslational modifications. In such cases other specialized programs such as ProMoST can be used.

Moreover, even if we consider that presented here new *pKa* values are different from those which were derived earlier experimentally, one should remember that even experimental setup can have strong impact on the results. For instance *pKa* values obtained by Thurkill et al. were measured using alanine pentapeptides with charged residue in the center. This was done to minimalize the contribution from neighboring residues, but from the other side, such a setup is extremely far from the real situation in the proteins (contribution from surrounding residues which are not Ala peptides, post translational modifications, etc.). Thus, optimized *pKa* values such as presented herein in some way are closer to the reality as they indirectly take into account such complexity. In the Supplementary Table 2 one can find average *pKa* values from previously used scales compared to IPC values. On peptide dataset most of differences is due terminal residues, which could be expected as in the peptides terminal charge can constitute big proportion of overall charge, thus N-terminus *pKa* value in previous studies was underestimated, while C-terminus *pKa* was overestimated in comparison to IPC values. On the other hand, for proteins one can notice that the main differences are observed for cysteine reflecting possible contribution from disulfide bridges and for lysine, histidine, and tyrosine which are frequently posttranslationally modified. Moreover, this effect is less abundant for arginine (also frequently modified), but it should be noted that arginine is bigger and contains more charged groups thus most likely modification effect (if exists) is less profound.

**Figure 3.**
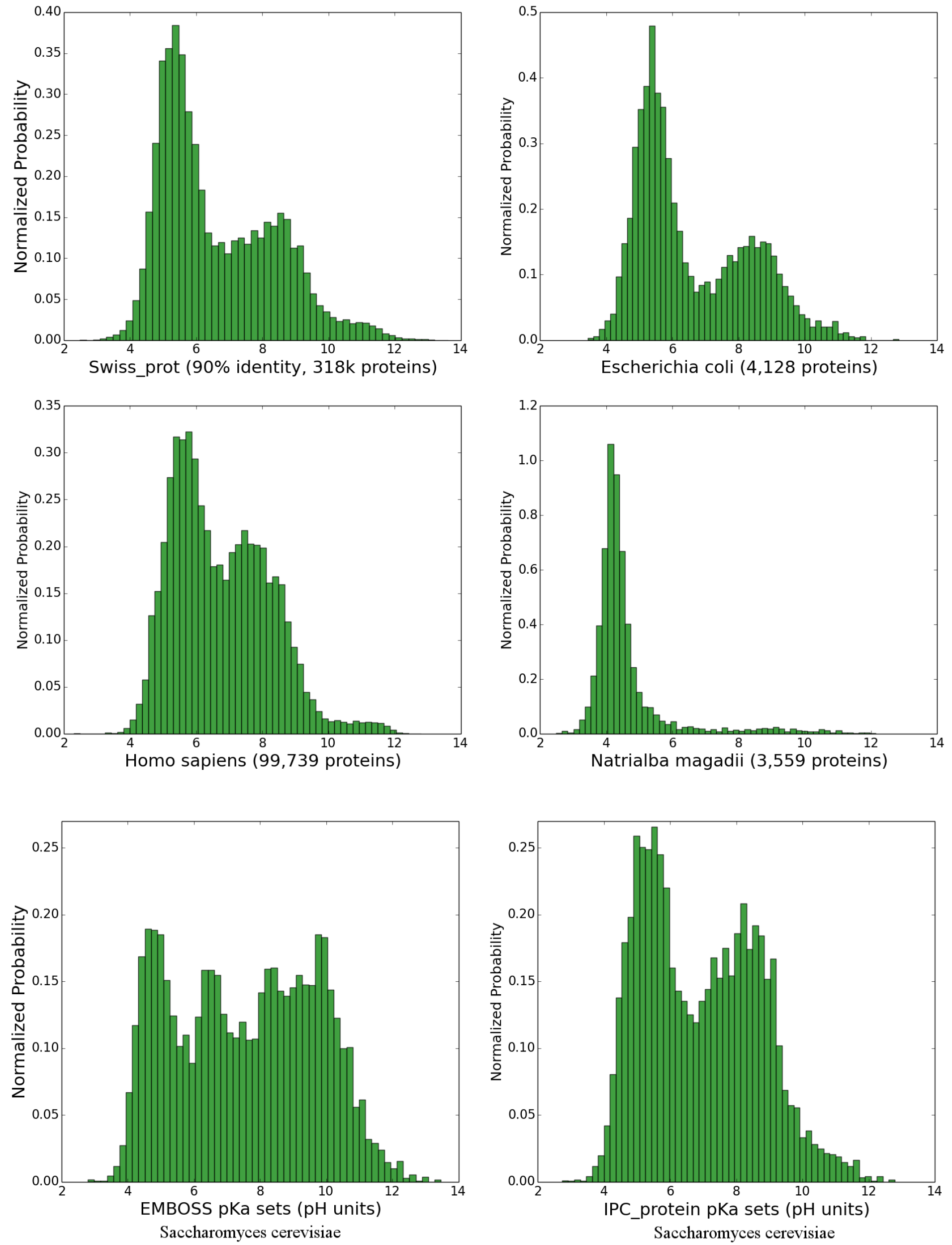
Histograms of the isoelectric points of proteins. Top and middle panels are calculated using the IPC_protein *pKa* set (in 0.25 pH unit intervals) and represents *pI* distribution in the *SwissProt* database, human proteome, *Escherichia coli* and extreme halophilic archaeon *Natrialba magadii*. Bottom two panels presents the isoelectric points of the yeast proteome (6,721 proteins) calculated using the EMBOSS *pKa* set (as presented in the Saccharomyces Genome Database14) and the IPC_protein *pKa* set for comparison.

## Methods

### Isoelectric point, Henderson-Hasselbalch equation, pKa values for the ionizable groups of proteins

The isoelectric point (*pI*) is the pH at which the net charge of a protein is zero. For polypeptides, the isoelectric point depends primarily on the dissociation constants (*pKa*) for the ionizable groups of seven charged amino acids: glutamate (δ-carboxyl group), aspartate (ß-carboxyl group), cysteine (thiol group), tyrosine (phenol group), histidine (imidazole side chains), lysine (ε-ammonium group) and arginine (guanidinium group). Moreover, the charge of the terminal groups (NH_2_ and COOH) can greatly affect the *pI* of short peptides. Generally, the Glu, Asp, Cys, and Tyr ionizable groups are uncharged below their *pKa* and negatively charged above their *pKa*. Similarly, the His, Lys, and Arg ionizable groups are positively charged below their *pKa* and uncharged above their pKa^5^. This has certain implications. For example, during electrophoresis, the direction of protein migration on the gel depends on the charge. If the buffer pH (and as a result, the gel pH) is higher than the protein isoelectric point, the particles will migrate to the anode (negative electrode), and if the buffer pH is lower than the isoelectric point, they will migrate to the cathode. When the gel pH and the protein isoelectric point are equal, the proteins stop to migrate.

Overall, the net charge of the protein or peptide is related to the solution (buffer) pH. We can use the Henderson-Hasselbalch equation^6^ to calculate the charge at a certain pH:

- for negatively charged macromolecules:

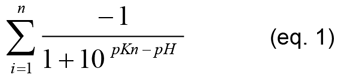

Where *pKn* is the acid dissociation constant of the negatively charged amino acid

- for positively charged macromolecules:

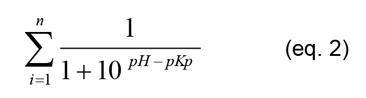

Where *pKp* is the acid dissociation constant of the positively charged amino acid

The charge of a macromolecule at a given pH is the sum of the positive and negative charges of the individual amino acids given by 1 and 2. When the *pKa* values are set, the only variable in the equations is the pH of the buffer, and by iteratively changing the pH, we can easily calculate the isoelectric point. The result will be almost certainly different than the real isoelectric point because many proteins are chemically modified (e.g., amino acids can be phosphorylated, methylated, acetylated), which can change their charge. The occurrence of cysteines (negative charge), which may oxidize and lose charge when they form disulfide bonds in the protein, is also problematic. Moreover, one must consider the charged residue exposure to solvent, dehydration (Born effect), charge-dipole interactions (hydrogen bonds), and charge-charge interactions^5^.

Nevertheless, the most critical consideration for accurate isoelectric point determination is the use of appropriate *pKa* values. Unfortunately, *pKa* estimates differ depending on the experimental setup in which they were measured. More than 600 different *pKa* values have been reported for the ionizable groups^15^. Table 4 shows the most commonly used values, including two new *pKa* sets (IPC_protein and IPC_peptide) proposed in this study. Most of the algorithms use nine parametric models (seven *pKa* values corresponding to charged amino acids and two for the terminal groups), but more advanced algorithms also exist, e.g., Bjellqvist16 (17 parameters) and ProMoST^17^ (72 parameters), which take advantage of specifying additional *pKa* values for charges of particular amino acids, especially those located on the polypeptide termini.

**Table 4.** Most commonly used *pKa* values for the ionizable groups of proteins. Note that Bjellqvist and ProMoST use different amounts of additional *pKa* values (not shown), which take into account the relative position of the ionized group (whether it is located on the N- or C- terminus or in the middle). For more details, see References 4 and 5 and the “Theory” section on the IPC web site.

| Amino acid | NH2 | COOH | C | D | E | H | K | R | Y |
| --- | --- | --- | --- | --- | --- | --- | --- | --- | --- |
| EMBOSS<sup>18</sup> | 8.6 | 3.6 | 8.5 | 3.9 | 4.1 | 6.5 | 10.8 | 12.5 | 10.1 |
| DTASelect<sup>19</sup> | 8 | 3.1 | 8.5 | 4.4 | 4.4 | 6.5 | 10 | 12 | 10 |
| Solomon<sup>20</sup> | 9.6 | 2.4 | 8.3 | 3.9 | 4.3 | 6 | 10.5 | 12.5 | 10.1 |
| Sillero<sup>21</sup> | 8.2 | 3.2 | 9 | 4 | 4.5 | 6.4 | 10.4 | 12 | 10 |
| Rodwell<sup>22</sup> | 8 | 3.1 | 8.33 | 3.68 | 4.25 | 6 | 11.5 | 11.5 | 10.07 |
| Patrickios<sup>23</sup> | 11.2 | 4.2 | - | 4.2 | 4.2 | - | 11.2 | 11.2 | - |
| Wikipedia | 8.2 | 3.65 | 8.18 | 3.9 | 4.07 | 6.04 | 10.54 | 12.48 | 10.46 |
| Lehninger<sup>24</sup> | 9.69 | 2.34 | 8.33 | 3.86 | 4.25 | 6 | 10.5 | 12.4 | 10 |
| Grimsley<sup>15</sup> | 7.7 | 3.3 | 6.8 | 3.5 | 4.2 | 6.6 | 10.5 | 12.04* | 10.3 |
| Toseland<sup>25</sup> | 8.71 | 3.19 | 6.87 | 3.6 | 4.29 | 6.33 | 10.45 | 12 | 9.61 |
| Thurlkill<sup>26</sup> | 8 | 3.67 | 8.55 | 3.67 | 4.25 | 6.54 | 10.4 | 12 | 9.84 |
| Nozaki<sup>27</sup> | 7.5 | 3.8 | 9.5 | 4 | 4.4 | 6.3 | 10.4 | 12 | 9.6 |
| Dawson<sup>28</sup> | 8.2** | 3.2** | 8.3 | 3.9 | 4.3 | 6 | 10.5 | 12 | 10.1 |
| Bjellqvist<sup>16</sup> | 7.5 | 3.55 | 9 | 4.05 | 4.45 | 5.98 | 10 | 12 | 10 |
| ProMoST<sup>17</sup> | 7.26 | 3.57 | 8.28 | 4.07 | 4.45 | 6.08 | 9.8 | 12.5 | 9.84 |
| **IPC_protein** | **9.094** | **2.869** | **7.555** | **3.872** | **4.412** | **5.637** | **9.052** | **11.84** | **10.85** |
| **IPC_peptide** | **9.564** | **2.383** | **8.297** | **3.887** | **4.317** | **6.018** | **10.517** | **12.503** | **10.071** |

\*Arg was not included in the study, and the average $pK_a$ from all other $pK_a$ sets was taken.
\*\* NH2 and COOH were not included in the study, and they were taken from Sillero.

## Datasets

The aim of the present study was to derive computationally more accurate *pKa* sets using currently available data. For training and validation, the following datasets were used:

- The IPC peptide *pKa* set was optimized using peptides from three, high-throughput experiments:

a. unmodified 5,758 peptides from Gauci et al.^29^ - peptides from zebrafish lysate fractionated using isoelectric focusing
b. PHENYX dataset (7,582 peptides)^4^ - peptides from Drosophila Kc167 cell line fractionated using isoelectric focusing on off-gel electrophoresis device
c. SEQUEST dataset (7,629 peptides)^4^ - peptides from Drosophila Kc167 cell line fractionated using isoelectric focusing on off-gel electrophoresis device

- The IPC protein *pKa* set was optimized using proteins from two databases:

a. SWISS-2DPAGE, release 19.2 (2,530 proteins)^30^ - based on literature data about *pI* linked to UNIPROT accession numbers
b. PIP-DB (4,947 entries)^31^ - based on literature data, provides *pI* and sequence information for about half of the records (for details see Table 5).

First, the raw data from the individual datasets were parsed to the unified fasta format with information about the isoelectric point stored in the headers. Next, datasets consisting of proteins and datasets consisting of peptides were merged into two datasets (IPC_protein and IPC_peptide, respectively). The data was carefully validated, e.g., if multiple experimental *pI* values were reported, the average was used. The first, major splicing form of the protein (most widely expressed) taken from UniProt^32^ was used for SWISS-2DPAGE. Additionally, the information about experimental methods used for obtaining isoelectric points or their specificity was not used implicitly during this study. Similarly, as the information about post translational modifications (PTMs) was included directly in SWISS-2DPAGE and PIP-DB, it was not possible to investigate PTMs contribution to *pI* and they were assumed to be absent. Outliers representing possible annotation errors in databases were removed (proteins with mean standard error (MSE) > 3 between the experimental isoelectric point and the average predicted *pI*; note that under this cutoff, no peptides were removed). Redundant data was removed using CD-HIT^33^ ( 0.99 sequence identity threshold was used; in this case, it was adequate to use such a high sequence identity because even single mutations in the charged residues can lead to dramatic changes in *pI*; moreover other sequence identity thresholds gave similar results; data not shown). This step also removed duplicates (multiple entries assigned to the same sequence coming from two different databases). Finally, 25% of the randomly chosen proteins and peptides were excluded for final testing, and the remaining 75% were used for 10-fold cross-validated training.

Detailed statistics for the datasets can be found in Table 5. All dataset files are available as Supplementary Files and/or online in the “Datasets” section of the IPC web site.

**Table 5.**
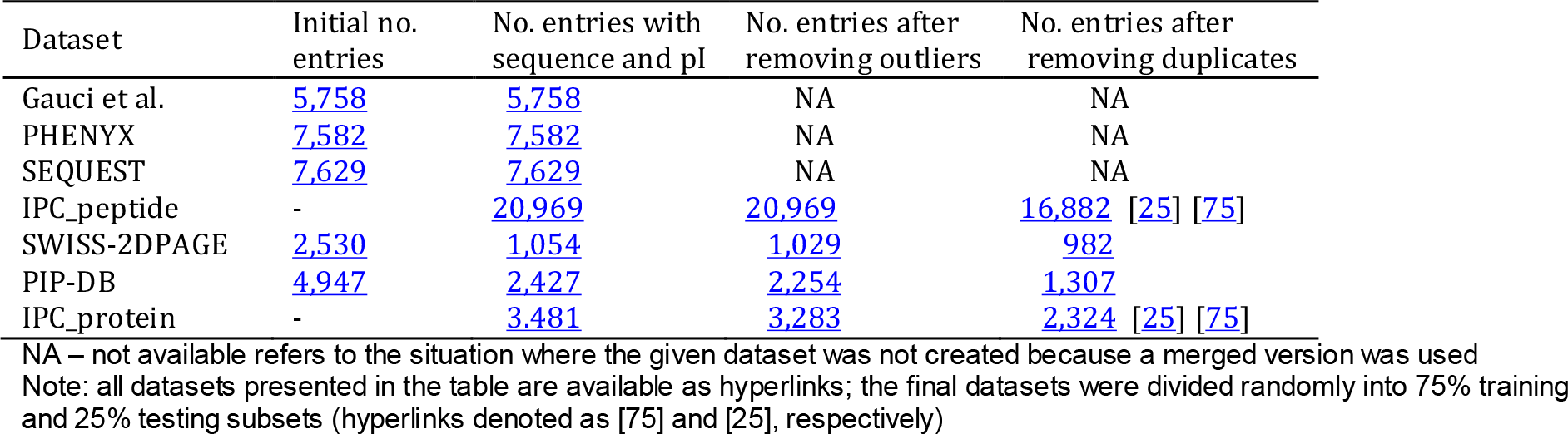
Detailed statistics for the available dataset

| Dataset | Initial no. entries | No. entries with sequence and *pI* | No. entries after removing outliers | No. entries after removing duplicates |
| --- | --- | --- | --- | --- |
| Gauci et al. | 5,758 | 5,758 | NA | NA |
| PHENYX | 7,582 | 7,582 | NA | NA |
| SEQUEST | 7,629 | 7,629 | NA | NA |
| IPC_peptide | - | 20,969 | 20,969 | 16,882 [25] [75] |
| SWISS-2DPAGE | 2,530 | 1,054 | 1,029 | 982 |
| PIP-DB | 4,947 | 2,427 | 2,254 | 1,307 |
| IPC_protein | - | 3,481 | 3,283 | 2,324 [25] [75] |

NA – not available refers to the situation where the given dataset was not created because a merged version was used
Note: all datasets presented in the table are available as hyperlinks; the final datasets were divided randomly into 75% training and 25% testing subsets (hyperlinks denoted as [75] and [25], respectively).

## Calculation of the isoelectric point

As noted before, the isoelectric point is determined by iteratively calculating the sum of Equations 1 and 2 for the individual charged groups for a given pH. The calculation can be performed exhaustively, but this would not be practical. Instead, the bisection algorithm^34^ is used, which in each iteration halves the search space (initially, the pH is set to 7) and then moves higher or lower by 3.5 (half of 7) depending on the charge. In the next iteration, the pH is changed by 1.75 (half of 3.5), and so on. This process is repeated until the algorithm reaches the desired precision. Bisection improves the speed by 3-4 orders of magnitude, and after approximately a dozen of iterations, the algorithm converges with 0.001 precision. Next, the speed improvement can be obtained by starting the search from a rough approximation of the solution rather than 7 (in this case, a pH of 6.68 was used, which is the average isoelectric point for approximately 318,000 proteins taken from the *SwissProt* database^35^, 90% sequence identity threshold was used).

## Performance measures

To measure the performance, two metrics were used i.e., the root-mean-square deviation (RMSD) and the number of outliers, defined as *pI* predictions with a mean standard error (MSE) larger than the given threshold in comparison with the experimental *pI*. To remove potential outliers, for the protein datasets, an MSE of3 was used, and for peptide datasets, an MSE of 0.25 was used. Moreover, for the preliminary analysis, the Pearson correlation was used.

## Optimization

The optimization procedure was designed to obtain nine optimal *pKa* values (corresponding to the N- and C-termini and the C, D, E, H, K, R, and Y charges). The cost function was defined as the root-mean-square deviation (RMSD) between the true isoelectric points from the available datasets and those calculated using the new *pKa* set(s). Optimization was performed using a Basin-Hopping procedure^10^ which uses a standard Monte Carlo algorithm with Metropolis criterion to decide whether to accept a new solution. The previously published *pKa* values were used as the initial seeds. To limit the search space, a truncated Newton algorithm^36^ was used, with 2 pH unit bounds for the *pKa* variables (e.g., if the starting point for Cys *pKa* was 8.5, the solution was allowed in the interval [6.5, 10.5]). The optimization was run iteratively multiple times using intermediate *pKa* sets until the algorithm converged and no better solutions could be found. To avoid overfitting, both the IPC_protein and IPC_peptide datasets were randomly divided into 75% training datasets (used for *pKa* optimization) and 25% testing datasets (not used during optimization). During training, nested 10-fold cross-validation was used^37^. Thus, the IPC was optimized separately on k-1 partitions and tested on the remaining partition. The training was repeated ten times in all combinations. The resulting *pKa* sets were averaged. In general, this process results in slower convergence of the algorithm and a longer training time but prevents overfitting. Apart from the nine parametric model (nine *pKa* values for charged residues) also more advanced models similar to Bjellqvist and ProMoST were also tested. Their performance was on a similar level thus the simpler, nine parametric model was used in the final version of IPC.

## Implementation

The IPC, Isoelectric Point Calculator is available as a web server (Fig. 4) implemented in PHP server-side scripting language. Additionally, HTML5 JavaScript charting library CanvasJS (http://canvasjs.com)and bootstrap (http://getbootstrap.com) were used. Moreover, IPC can be used on any operating system as a standalone program written in Python language (Supplementary File 3).

**Figure 4.**
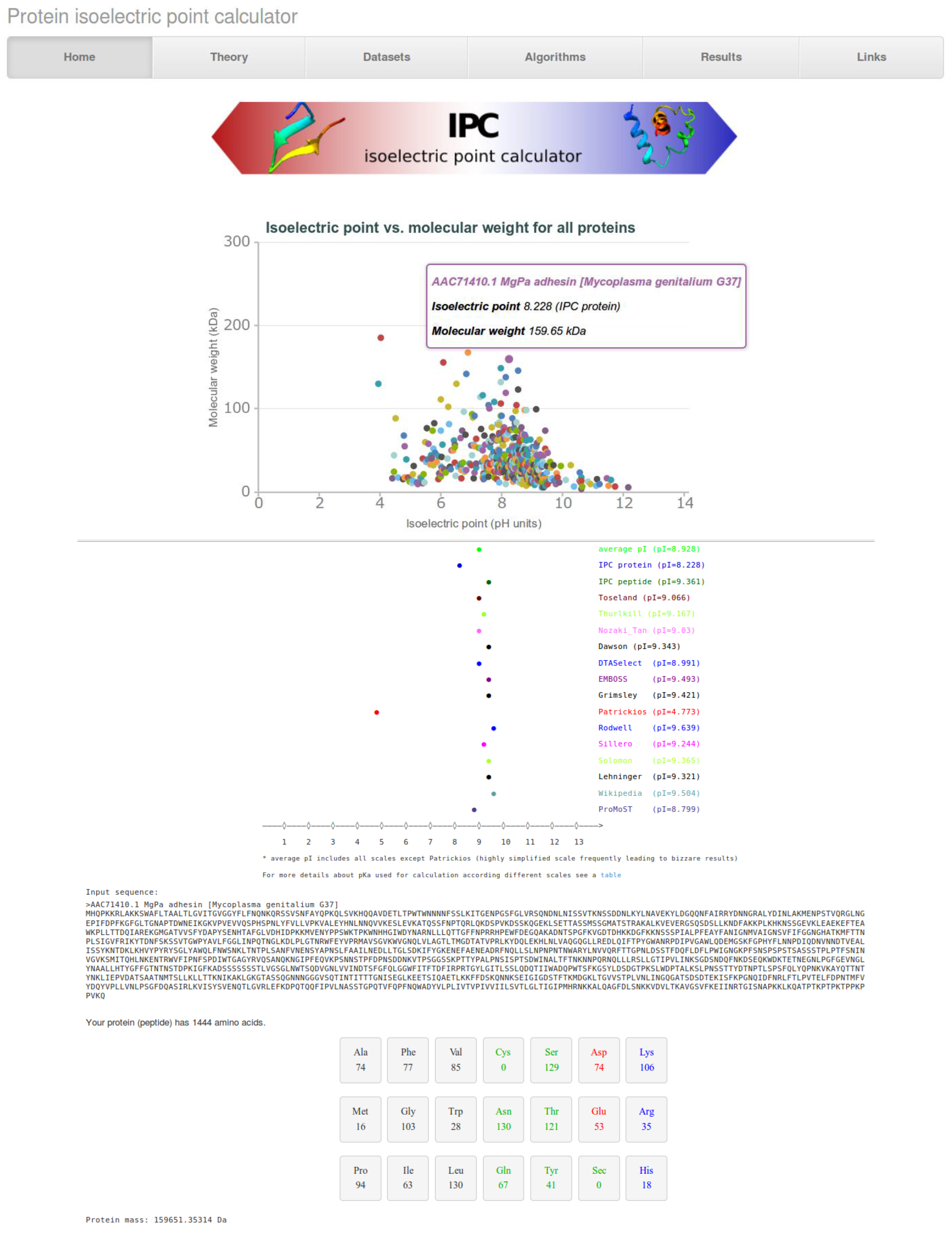
Exemplary output of the IPC calculator for the *Mycoplasma genitalium* G37 proteome (a highly reduced organism with 476 proteins). The scatter plot with the predicted isoelectric points versus molecular weight for all proteins is presented at the top. Then, for individual proteins, *pI* predictions based on different *pKa* sets are presented alongside the molecular weight and amino acid composition.

## Acknowledgements

LPK acknowledges all authors of previous works related to different *pKa* sets and datasets, especially developers of SWISS-2DPAGE database. The author thanks also Yasset Perez-Riverol for assistance with pIR package and Vladlen Skvortsov for assistance with pIPredict program. Additionally, LPK would like to thank all members of the Soeding lab for fruitful discussions. Last but not least, LPK acknowledge useful suggestions from anonymous reviewers.

## Author contributions statement

LPK conceived and developed the study, analyzed and interpreted the experiments, and wrote the article.

## Supplementary Information

- Supplementary file 1 - IPC peptide dataset (16,882 peptides, derived from Gauci et al. PHENYX and SEQUEST after 99% redundancy removal) – fasta formattedisoelectric.ovh.org/datasets/GauciPHENYXSEQUEST0.99duplicatesout.fasta.doc.zip
- Supplementary file 1-IPC peptide dataset (16,882 peptides, derived from Gauci et al.PHENYX and SEQUEST after 99% redundancy removal)-fasta formatted isoelectric.ovh.org/datasets/GauciPHENYXSEQUEST0.99duplicatesout.fasta.doc.zip
- Supplementary file 2 - IPC protein dataset (2,324 proteins, derived from SWISS-2DPAGE and PIP-DB after 99%redundancy removal)-fasta formattedisoelectric.ovh.org/datasets/pipch2d1921stisoformoutliers3unitscleaned0.99.fasta.doc.zip
- Supplementary file 3 - IPC Isoelectric Point Calculator source code (python, any OS)isoelectric.ovh.org/IPCstandaloneversion.zip Moreover, as stated in the manuscript all datasets combinations used in study are available as hyperlinks in Table 5
(22 additional files) and online isoelectric.ovh.org/datasets.html

**Table S1.**
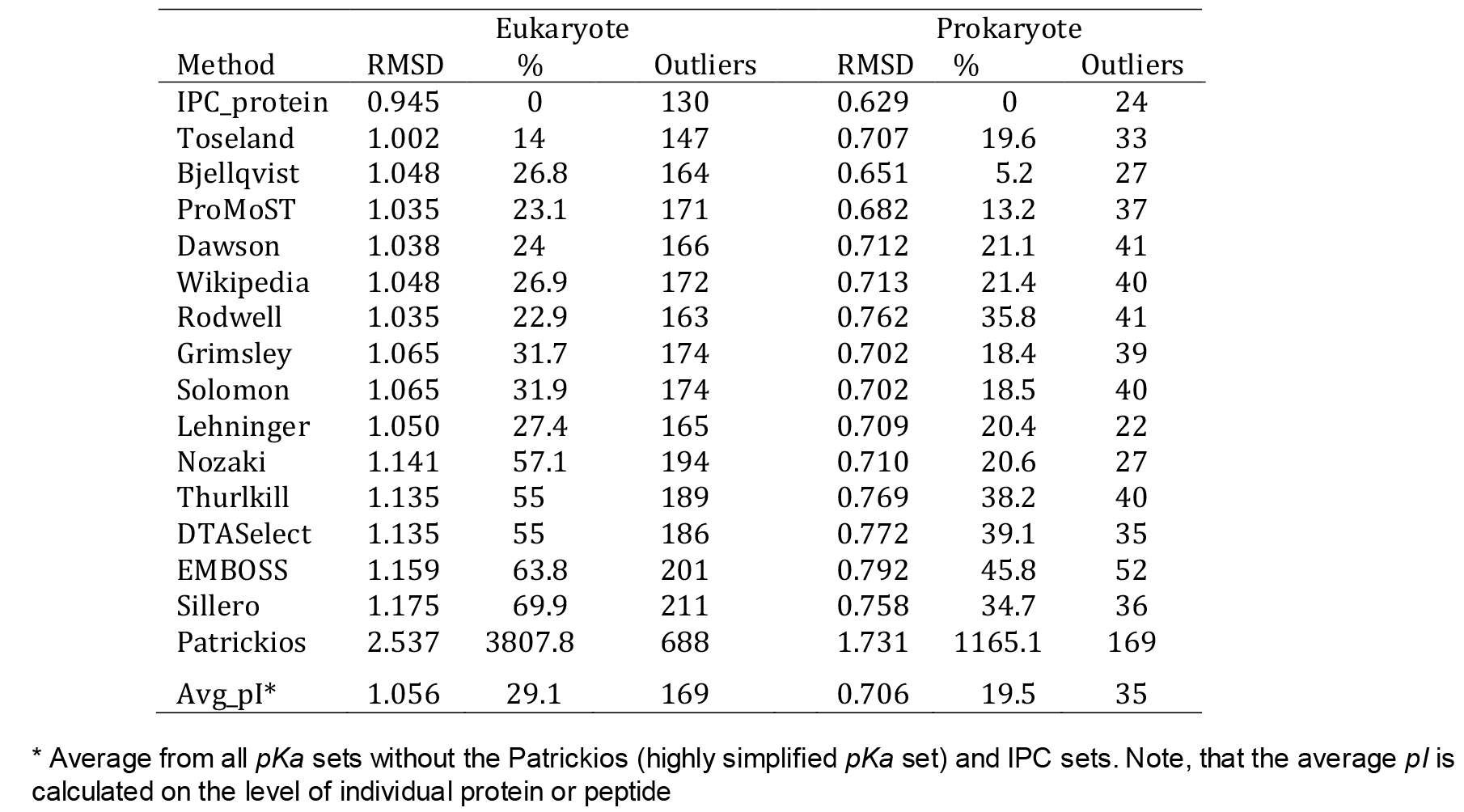
Performance of isoelectric point prediction algorithms on prokaryotic (837 proteins) and eukaryotic (1455 proteins) datasets derived from SWISS-2DPAGE and PIP-DB.

**Table S2.**
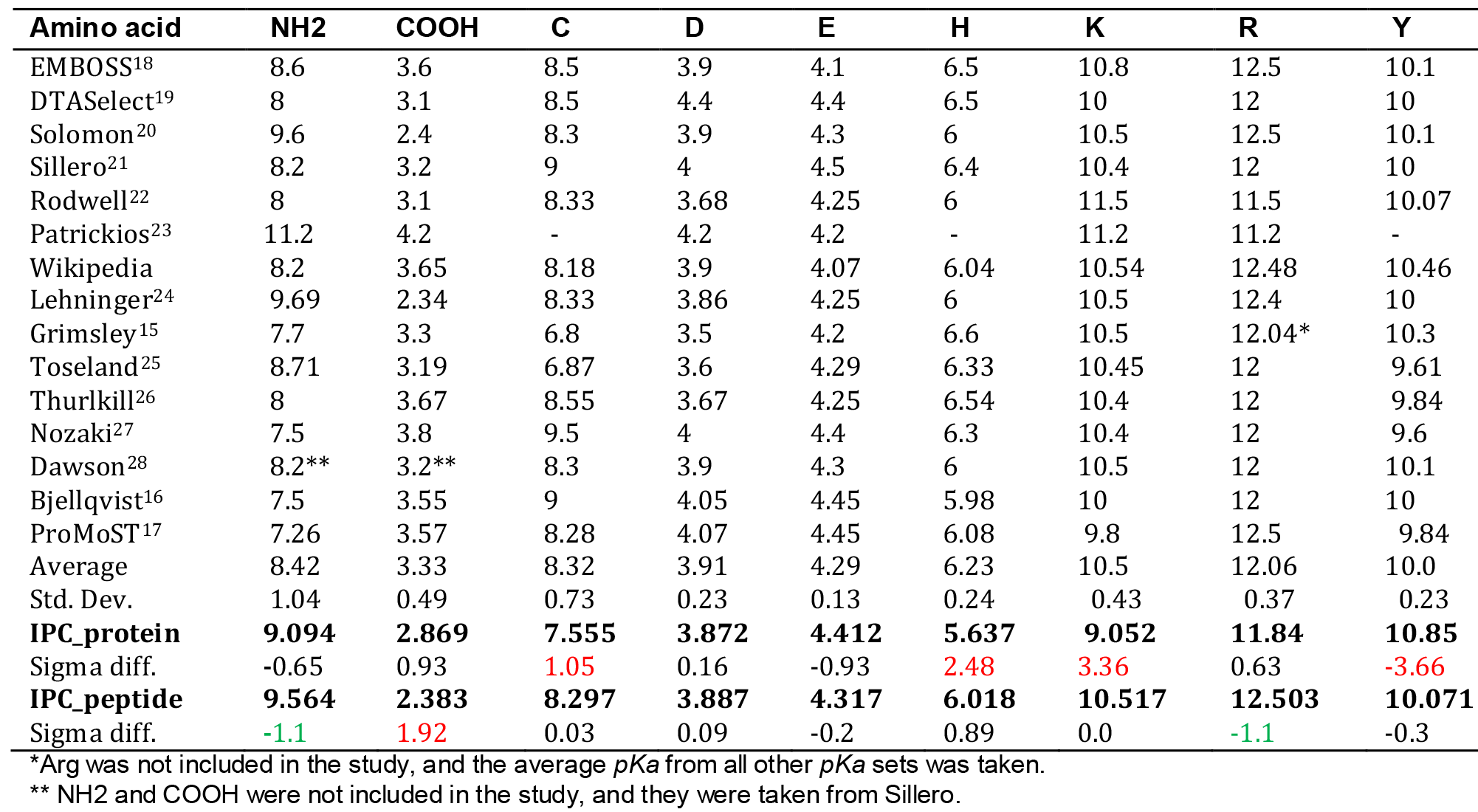
Statistical comparison of previous pKa values to IPC_protein and IPC_peptide sets. The differences bigger than one sigma were marked in red (positive) and green(negative)

| Amino acid | NH2 | COOH | C | D | E | H | K | R | Y |
| --- | --- | --- | --- | --- | --- | --- | --- | --- | --- |
| EMBOSS<sup>18</sup> | 8.6 | 3.6 | 8.5 | 3.9 | 4.1 | 6.5 | 10.8 | 12.5 | 10.1 |
| DTASelect<sup>19</sup> | 8 | 3.1 | 8.5 | 4.4 | 4.4 | 6.5 | 10 | 12 | 10 |
| Solomon<sup>20</sup> | 9.6 | 2.4 | 8.3 | 3.9 | 4.3 | 6 | 10.5 | 12.5 | 10.1 |
| Sillero<sup>21</sup> | 8.2 | 3.2 | 9 | 4 | 4.5 | 6.4 | 10.4 | 12 | 10 |
| Rodwell<sup>22</sup> | 8 | 3.1 | 8.33 | 3.68 | 4.25 | 6 | 11.5 | 11.5 | 10.07 |
| Patrickios<sup>23</sup> | 11.2 | 4.2 | - | 4.2 | 4.2 | - | 11.2 | 11.2 | - |
| Wikipedia | 8.2 | 3.65 | 8.18 | 3.9 | 4.07 | 6.04 | 10.54 | 12.48 | 10.46 |
| Lehninger<sup>24</sup> | 9.69 | 2.34 | 8.33 | 3.86 | 4.25 | 6 | 10.5 | 12.4 | 10 |
| Grimsley<sup>15</sup> | 7.7 | 3.3 | 6.8 | 3.5 | 4.2 | 6.6 | 10.5 | 12.04* | 10.3 |
| Toseland<sup>25</sup> | 8.71 | 3.19 | 6.87 | 3.6 | 4.29 | 6.33 | 10.45 | 12 | 9.61 |
| Thurlkill<sup>26</sup> | 8 | 3.67 | 8.55 | 3.67 | 4.25 | 6.54 | 10.4 | 12 | 9.84 |
| Nozaki<sup>27</sup> | 7.5 | 3.8 | 9.5 | 4 | 4.4 | 6.3 | 10.4 | 12 | 9.6 |
| Dawson<sup>28</sup> | 8.2** | 3.2** | 8.3 | 3.9 | 4.3 | 6 | 10.5 | 12 | 10.1 |
| Bjellqvist<sup>16</sup> | 7.5 | 3.55 | 9 | 4.05 | 4.45 | 5.98 | 10 | 12 | 10 |
| ProMoST<sup>17</sup> | 7.26 | 3.57 | 8.28 | 4.07 | 4.45 | 6.08 | 9.8 | 12.5 | 9.84 |
| Average | 8.42 | 3.33 | 8.32 | 3.91 | 4.29 | 6.23 | 10.5 | 12.06 | 10.0 |
| Std. Dev. | 1.04 | 0.49 | 0.73 | 0.23 | 0.13 | 0.24 | 0.43 | 0.37 | 0.23 |
| **IPC_protein** | **9.094** | **2.869** | **7.555** | **3.872** | **4.412** | **5.637** | **9.052** | **11.84** | **10.85** |
| Sigma diff. | -0.65 | 0.93 | 1.05 | 0.16 | -0.93 | 2.48 | 3.36 | 0.63 | -3.66 |
| **IPC_peptide** | **9.564** | **2.383** | **8.297** | **3.887** | **4.317** | **6.018** | **10.517** | **12.503** | **10.071** |
| Sigma diff. | -1.1 | 1.92 | 0.03 | 0.09 | -0.2 | 0.89 | 0.0 | -1.1 | -0.3 |

\*Arg was not included in the study, and the average $pK_a$ from all other $pK_a$ sets was taken.
\*\* NH2 and COOH were not included in the study, and they were taken from Sillero.

